# Effect of landscape structure on genetic structure of the Lesser horseshoe bat (*Rhinolophus hipposideros*) in Britanny colonies

**DOI:** 10.1101/2021.11.26.470144

**Authors:** Alexandra Rodriguez, Eric Petit

## Abstract

Some species are difficult to observe and others, need to be not disturbed because of their vulnerability. In response to the difficulty of studying the dispersal behaviors of these species, some areas of biology have been combined in order to access the information despite practical limitations. Here we present the combination of several methodologies from landscape ecology to non-invasive population genetics that allow us to obtain important information on *Rinolophus hipposideros*, a vulnerable European bat. We genotyped 18 georeferrenced colonies in Brittany (France) from droppings collected in their refuges. We used 6 microsatellite markers in order to obtain the genetic distances between them. On the other hand we calculated Euclidian distances between the refuges occupied by these colonies and some ecological distances with the Pathmatrix module of ArcGis 3.2. We tested hypothesis about the difficulty of dispersal of the species in areas without forest cover or with a low density of hedges. Thanks to the Monmonier algorithm we could infer possible genetic barriers between the colonies and we could compare their location to the presence of landscape barriers (areas with little tree cover). We detected a pattern of isolation by distance that reveals limited dispersal capacities in the species but no pattern linked to ecological distances. We found that some of the neighboring colonies with greater genetic distances between them were located in areas with low density of hedges which could suggest an impact of this landscape element in their movements. Finer studies should allow us to conclude on the need or not of forest cover in the dispersal of this species.

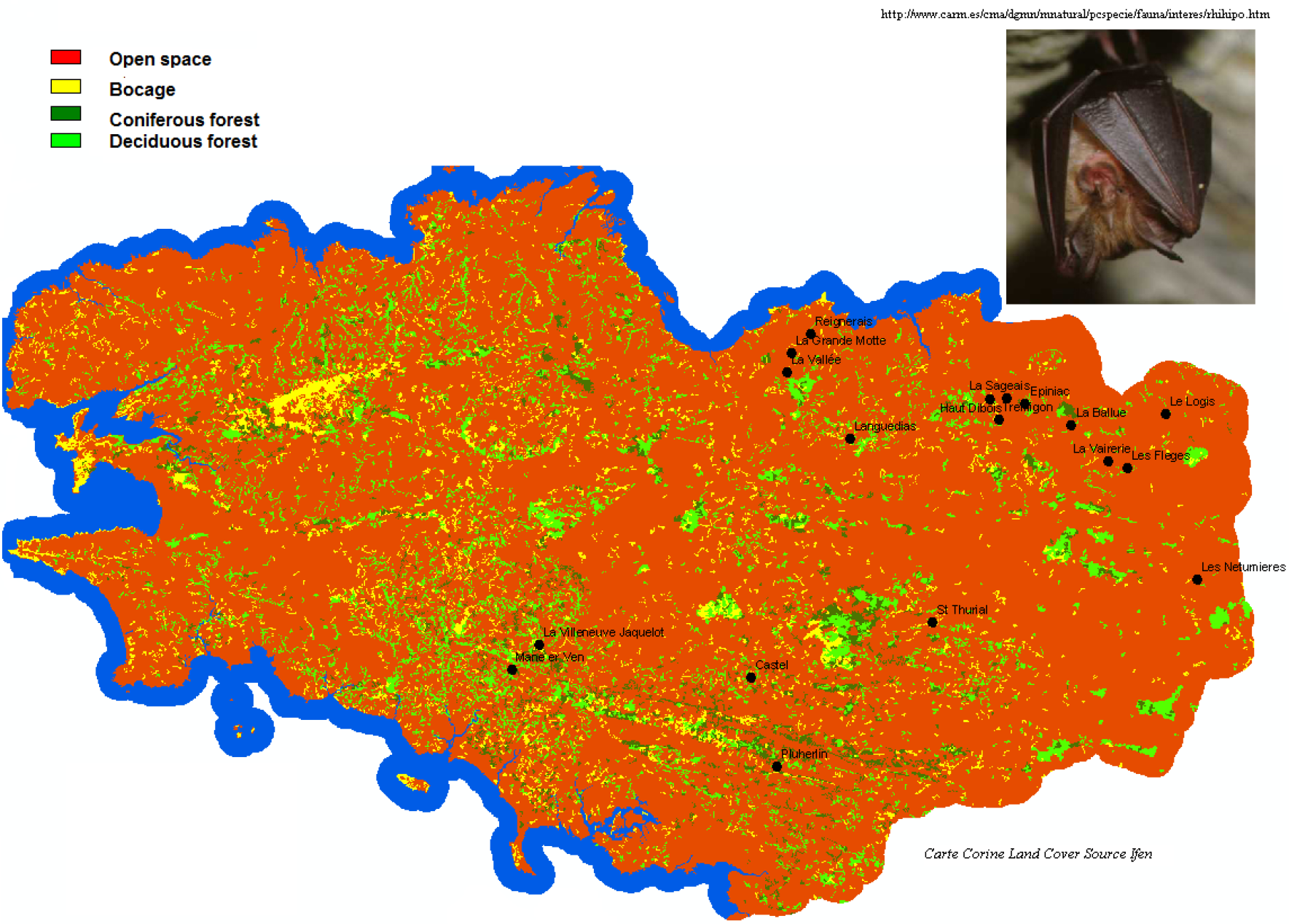

## Introduction

Since the Rio Convention in 1992, several processes have been highlighted as mechanisms having strong impacts on biodiversity. Habitat fragmentation is the process that has the stronger impact on animal species decline (Fischer and Lindenmayer 2002). During the last fifty years, landscape ecologists have studied the effects of habitat structure on animal populations’ structure and dynamics using biodiversity indices (Burel et Baudry 1999).

More recently, and inheriting some concepts and methods from landscape ecology a new discipline has emerged that links this area with population genetics. This hybrid discipline, landscape genetics, allows us to determine what are the landscape features that contribute to the genetic evolution of populations by their permeability or their difficulty to be crossed (Manel et al. 2003; Coulon et al. 2004; Funk et al. 2005). These aspects are developed in percolation theory (Stauffer and Aharony 1985).

Studies on population genetics use more and more frequently the available information on soil use and occupancy and the existent SIG tools that facilitate the description and quantification of landscape elements in order to involve them in biological processes analysis.

Several types of approaches are developed today and they are very useful to better understand microevolutive phenomena.

A first kind of method called “a priori” consists in testing directly the potential correlations that can exist between genetic distances of pairs of populations and habitat structure. In this case, habitat structure is represented by ecological distances that are the distances of the less costly paths existing between populations. The way to comp ute these distances consists in inferring costs of movement taking into account the animals difficulties to cross each type of habitat. In this kind of approach, the cost of crossing a landscape feature can be deduced from the knowledge on species’ anatomical, physiological and behavioral aptitudes (Keyghobadi et al. 2005 ; Poissant et al. 2005 ; Spear et al. 2005 ; Broquet et al. 2006a, 2006b ; Stevens et al. 2006).

A second type of method, called « a posteriori » consists in starting by making an analysis of genetic data without taking landscape structure into account. Only after this step we look if there are landscape features that can explain the patterns observed by genetic analysis. Two types of methods can be used for this purpose depending on the kind of data that we have. If we study discrete populations clearly defined and delimited in space we can infer genetic barriers existence between groups of populations using Monmonier algorithm because it allows us to separate the most genetically divergent groups (Manni et al.2004). On the contrary, when we have individuals that are dispersed on a surface in a continuous way and we do not know the extension of each population we can use Bayesian methods to assign each individual to a population by computing and weigh their probability to belong to each population. In this case we use the geographical coordinates of the individuals and genetic barriers are interpolated by kriging tools (Pritchard et al. 2000 ; Fallush et al. 2003 ; Guillot et al. 2004, 2005a, 2005b). Once we have found the genetic barriers we compare their loca tion with the presence of landscape features in order to know if they correspond to the presence of ecological barriers (rivers, mountains, etc). In this study we used both an “a priori” and an “ a posteriori” method in the analysis of the potential effect of landscape structure on genetic structure of 18 colonies of *Rhinolophus hipposideros* in Brittany region (France). This bat species has disappeared in Germany and Belgium during the last 20 years and has been declining at European scale. It has been classified several times as a vulnerable species by the IUCN. By consequence it seems important to understand the dynamics that may explain this decline and in particular the factors that are involved in this phenomenon in order to develop pertinent conservation projects. There are several hypotheses that try to explain the decline of *Rhinolophus hipposideros* (Bontadina et al. 2002). Some of these hypotheses are: the habitat fragmentation, the decrease of insect resources as dipters, trichopters and lepidopters that are essential to the Lesser horseshoe bat which is an insectivorous mammal. There is also an hypothesis concerning a direct toxic effect of insecticides on the wood of constructions and the decline of suitable sites by ancient buildings destruction in rural areas (Cosson et al. 2003).

The aim of this study was to test the hypothesis of habitat fragmentation. Several studies conducted on the field have found that *Rhinolophus hipposideros* systematically uses hedges when he goes hunting in the woods.

These data are true at local fine scale; the question here is if at a bigger scale the Lesser horseshoe bat displays these kinds of behavioral patterns. If this is the case, we can think that the suppression of forests and hedges could lead to an isolation of populations. We used8 microsatellite markers and maps of soil occupation in Brittany to conduct the study.

## Material and methods

### Rhinolophus hipposideros

The Lesser horseshoe bat , *Rhinolophus hipposideros,* is the smaller European bat species, with a weight comprised between 4 and 9g, with wingspan comprised between 192 nd 254mm. The species is present in occidental, meridional and central Europe (from western Ireland and Poland to southern Maghreb and from Atlantic Ocean to Danube Delta (Natura 2000). *Rhinolophus hipposideros* uses natural or artificial cavities (galleries and mines, pits, cellars, tunnels, military bunkers, caves) as hibernation spots. These locations are frequently underground and have precise characteristics: total obscurity, temperature comprised between 4 and 16°C, high values of hygrometry and total quiet. Concerning the birth spots, *Rhinolophus hipposideros* uses principally the top of buildings and caves (farms, churches, windmill) hot and clear (Holzhaider et al. 2002). This is an insectivore species and its diet composition varies with seasons and insect richness. The principal groups that *Rhinolophus hipposideros* consume are: dipters, lepidopters, nevropters and tricopters associated with humid woods (Natura 2000; Roué et al 1999). The hunting behavior associated with this insectivore diet presents several important steps. Preys’ capture is almost always realized during the purchase against leaves and on the ground, sometimes from a perch (Shoelfield et al. 2003). The individuals generally hunt alone in woody humid environments and less frequently in groups in open spaces where insects gather close to bovine excrements (McAney et Fairley 1988 ; Jones et Rayner 1989 ; Schofield 1996 ; Motte et Libois 2002). Moreover, many radio-tracking and bioacoustic studies based on ultrasonic sounds led to define precisely the foraging behavior of *Rhinolophus hipposideros* at a fine scale. Foraging takes place in woody areas, essentially in deciduous trees rich in insects and close from humid zones. Actually, specific studies on females’ behavior found a differential use of habitats. For example, a telemetric study conducted in Germany has demonstrated that females spent 39% of their foraging time in wood ensembles containing more than 50% of beeches whereas they spent only 34% of their time in mixed forests containing beeches and firs (15% of their time in forests with more than 50% of firs and 12% of their time in clearings (Holzhaider et al. 2002). Other studies have been able to make a classification of habitats in a decreasing preferential order for foraging: dense vegetation (aubepine and noisetiers)>deciduous forest (chenes, frenes, hetres, cornouillers)>mixed forests> farms with water points> hedges>coniferous forests (pins, sapins)>prairies (McAney et Fairley 1988 ; Scholfield et al. 1996, 2003 ; Tibbels et Kurta 2003). Moreover, it is important to note that Rhinolophus hipposideroas has been systematically observed hunting in hedges and using them to join feeding points. Some authors precise that individuals leave the caves by grillages, doors, and windows; they would follow the hedges or the walls and the roofs when vegetation is poor. Then they would use tunnels and bridges avoiding roads (Scholfield et al. 2003).

Knowing that the Lesser horshoe bat is highly predated by Pelrim falcon, Falcus peregrines and by Tyto alba; Motte and Libois (2002) made the hypothesis that this behaviors (hunting alone in wooded areas at less than 1m from the hedges and hunting in groups should decrease the predation risks. Moeover, othe authors that have classified the morphological and echolocation characteristics of the chiropters relating ir with species ecology conclude that there were two types of bats. There is a group of bat species with long and thin wings not very manœuvrable but conferring a rapid flight that will be more able to fly long distances in oen spaces. In the other hand there would be bats with short large wings highly manoeuvrable. These kind of wings allow flight in highly wooded areas but they are limited for movements outside of the forests (Norberg et Rayner 1987 ; Neuweiler 1989, 1990 ; Rhodes 1995 ; Bontadina et al. 2002).

In the Norberg and Rayner classification, Rhinolophus hipposideros is the Rhinolophidae with the smaller ration of length/width, so he is the most adapted to foraging in high density woody covert. Concerning the echolocation system, the produce ultrasonic sounds of constant frequency at 112kHz wtith a big cycle that allow them to detect the insects that are in movement in the air. This is probably the condition for having a selective foraging behavior (Neuweiler 1990 ; Bontadina et al. 2002). The high frequency is useful to detect little objects but it is very rapidly attenuated and I makes more difficult the detection of objects at big distances (Jones et Rayner 1989 ; Schnitzler et al. 2003). These two specializations (morphological and echolocation one) may be considered as adaptations to high density vegetation foraging. *Rhinolophus hipposideros* whould have evolved in this type of environment and the suppression of woody elements in the landscape could endangered its survival because he is not adapted to open spaces. Finally, several authors have demonstrated that Rhinolophus hipposideros was not used to travel big distances. In their ecological model underlined by the rate length of wings/width of wings related to ecological conditions, Norbergand Rayner (1987) predicted a foraging area comprised in a radius of 1.3km around the refuge for *Rhinolophus hipposideros*. Radio-traking studies have demonstrated that females that were not breastfeeding could forage in a radius comprised between 1,9 and 4,1km around the refuges (Bontadina et al. 2002), whereas pregnant females use less extended areas and stay at less than 600m from the refuge. Harmata has identifyied several types of movement in Rhinolophus hipposideros : frequent flights from 1 to 5km between the different seasonal refugees, rare migrations of more than 10km and a record of 146km between an summer refuge and an hibernation refuge (Harmata1989). He did not detect any tendency concerning short migrations (in all the directions) but long migrations tended to be conducted in a north-west south-east direction.

The previous studies conducted on Rhinolophus hipposideros that we mentioned before have shown us how this species uses its habitat at a small and proximal scale. If big scale dispersal follows the same patterns that the species displays during foraging behavior we could formulate the following hipotheses:

– *Rhinolophus hipposideros* can not travel big distances, migration and genetic exchange take place in a stepping stone landscape following Japanese steps inducing an isolation by distance (that could be observed by the increasing of genetic distance when euclidian distance increases between colonies)
– *Rhinolophus hipposideros* uses big forests and hedges when he travels. The colonies separated by an absence of these kinds of landscape features would be genetically more divergent in this case

### The samplings

We studied 18 colonies of the 50 known colonies in Brittany. For this study we worked on DNA from excrements collected from 18 Brittany colonies. The 18 colonies where chosen in three departments where *Rhinolophus hipposideros* is present (Ille et Vilaine, Côtes d’Armor and Morbihan). These colonies are separated by woody areas and by open spaces. Consequently, this structure should allow us to determine if they have an impact on population genetics. The study has been conducted on 540 bat droppings (36 samplings for each of the 15 colonies). Droppings were collected in the refuges with journal paper placed on the ground and conserved individually in 1,5ml test tubes with a piece of silica in order to absorb moisture and to avoid DNA degradation.

### DNA extraction

DNA extraction was conducted following the protocol developed by Mathy (2004) for bat droppings.

Extraction Protocol using DNeasyStool K it : Before starting you have to heat the oven to 70°C

If ASL buffer precipitates you have to put it in the steamroom at 70°C in order to dissolve it.

- Put the droppings individually in 2ml tubes, ad 1.6 ml of ASL buffer, crush the droppings with a toothpick as finely as possible
- Vortex each sampling during 1 minute (lysis and dissolution of the DNA)
- Ad an INHIBITEX® tablet in each tube et vortex during 1 minute (fixation of PCR inhibitors)
- Centrifuge the samplings at 13200 rpm during 6 min (proceed by groups of six samplings each time because the precipitate is easily resuspended)
- Prepare 25 μl of k proteinase in 2 ml tubes
- When centrifugation stops, take 600 μl of supernatant with a pipet in the tubes that contain k proteinase
- Ad AL buffer (600 μl) (do not mix AL buffer with k proteinase because it loses its effect) and incubate during 15minutes at 70°C
- Ad iced isopropanol (600 μl) and mix the tubes in order to homogenize the solution
- Number the columns Q iamp
- Transfer the samplings 600 by 600 μl on the columns and centrifuge during 1 min at 7200 rpm
- Repeat this operation until all the solution has been transferred to the column
- Ad 500 μl of buffer AW1 (firts wash) and centrifugate 1 min at 7200 rpm
- Ad 500 μl of buffer AW2 (second whash) and centrifugate 1 min à 7200 rpm
- Put the column on the numbered tubes
- DNA in 80μl of pure water
- Incubate 5 minutes at ambient temperature
- Centrifugate 1 minute at 7200 rpm

A series of extractions was conducted by colony (36 extractions and a white witness in order to detect potential contaminations).

To verify DNA presence in the extractions, each sampling was tested with a mitochondrial marker. We used Cytochrome b marker containing 350 pairs of bases. This marker was amplified by PCR with 1μl of DNA extract, 0,4 μl de of each primer, a volume of Taq buffer 0,6 units of Taq polymerase and 0,16mM of dNTPs.

Amplification was conducted in a thermal cycler PTC-100 (MJ Research) in the following conditions: DNA denaturation at 94°C during 15min, 50 cycles of 30s at 95°C, 30s at 55°C, 30s at 72°C and 10min for extension at 72°C. PCR products were colored with bromide staining and revealed by an ultraviolet light after 45 min of migration in an agar gel (120 V, 90mA) at 2% containing ethidium bromid. Depending on the band intensity, each sampling was classified into: negative (band absence), weak (band of low intensity) or positive (high intensity band).

### Genotypes sampling

We used an identification protocol based on eight microsatellite markers (D2, D119,C108, D111, D9, D113, D102, D103) amplified with a multiplex PCR. We used Puechmaille et al. (2005) protocol where 1 μl of sampling is mixed with 7 μl of Master mix (Multiplex, Qiagen) and 16 primers corresponding to the 8 markers. For each pair of primers one of them was marked with a fluorochrome allowing the analysis with the sequencer. The amplifications were conducted in a thermocycler PTC-100 in the following conditions : 15min at 95°C, 45 cycles at 94°C during 45s, 56°C during 45s, 72°C during 1min, 1 hour at 72°C.

Genotypes determination was done with two sequencers ABI PRISM 3130 XL Genetic Analyzer 16 Capillary system (Applied Biosystems) and with 500 Liz marker. For each sampling, 1 μl of PCR product was diluted in 3 μl ultrapure water. 1 μl of this mix was extracted and mixed with 0,5 μl of 500 Liz and 8,5 μl of formamide.

Genotypes lecture was conducted with GeneScan and Genemapper. Each sampling was amplified two times independently and a consensual genotype was elaborated for each locus with the two genotypes obtained during the amplifications. The protocol was repeated using Puechmaille and collaborators method (2005) until obtaining the consensual genotype for a maximum of samplings and a maximum of loci. The aim of this method was to identify the different samplings and to determinate which samplings corresponded to the droppings of a same individual in order to use only one copy when computing the allelic frequencies. The identification of samplings corresponding to the same individual was conducted with GeneCap software (Wilberg and Dreher 2004) which allows the comparison of obtained genotypes pair by pair.

### Genetic data analysis

Before analysing the impact of habitat structure on genetic structure we analyzed populations’ characteristics.

First of all we tested if they were at Hardy Weinberg equilibrium and if there was an allelic disruption or not. Differences from Hardy Weinberg equilibrium may be explained either by genotyping errors or by a selection pressure on a particular locus or by non aleatory reproduction between individuals in a same group (Sacks et al. 2004). Genetic diversity was characterized with two indices, allelic richness (El Mousadik & Petit 1996) and expected heterozygosity.

The whole study is based on the analysis of eventual similarities or differences between colonies. We distinguish two methods to compute them. When we work on discrete populations clearly defined in space we can use Fst values between populations in order to compute distances between them. Genetic distance is obtained by Fst/(1-Fst) formula (Rousset 1997). On the contrary, when populations are distributed in a continuous way on the studied area, we work using genetic the distance ar = (Qw-Qr)/(1-Qw) where Qr is the probability of identity of genes taken from two individuals separated by geographical distance (r) and where Qw is the probability of having two identical genes in the same individual (Rousset 2000).

In the present case we deal with colonies that belong to particular refuges, thus to discrete sets, that is why we used Fst for computing genetic distances. We conducted these analyses with the Fstat software (Goudet 1995).

### Analysis of the impact of landscape structure on genetic structure using an « a priori » method

The « a priori » analysis consists in testing several explicative models on populations’ genetic differentiation using different hypothesis concerning landscape use by animals. In this approach, we start from the postulate that an animal is not confronted to the same movement difficulties in the different habitats of a heterogeneous landscape (Burel et Baudry 1999). The idea is by consequence based on the trial to encode the landscape features with travelling costs that are a priori defined by data concerning the species’ biology. Once the different elements of the landscape are defined in this context and for a particular hypothesis, we look for the less costly paths between pairs of colonies. Finally, we test the correlation between these ecological distances (cost or distance of the less costly path) and the genetic distances. We repeated this operation for the different hypothesis in order to found which model explains better the genetic structure of populations.

For the application of this « a priori » method we computed the ecological distances with software ArcGIS 3.2 and its package Pathmatrix developped by Ray (2005). In order to be able to compute these distances, the colonies were previously georeferenced on Britanny map. Georeferencing has been done with aerial photographies and IGN departmental maps on carto Explorer software with Extended Lambert II coordinates. We used the ArcGIS 3.2 extension based on an algorithm which simulates the different possible paths between colonies. This algorithm computes their cost, it indentifies the less costly path and it gives it length and it cost in the feedback result.

We realized several necessary steps for the obtention of the ecological distances. First of all we used a Corine Land Cover map from Ifen (Institut français pour l’environnement) which represents the Brittain soil in 2000. On this map we can count 34 types of soil occupancy represented by adjacent polygons of 25ha pixels. These 34 types were classified with Arc GIS software in four pertinent and simple groups for the study: open spaces, deciduous forest, coniferous forest and bocage (terrain of mixed woodland and pasture with hedges). Then, each one of these soil types was affected with a travel cost. Finally, we rasterized the map in pixels of 25ha. It is necessary to use the raster version of the map for the real sum of ecological distances between colonies as it allows to compute all the pixels comprised between the colonies and to sum the travel cost within each of them. This method was used six times (once by each tested hypothesis). We tested three different types of hypothesis. The first one was the hypothesis dealing with isolation by distance in order to have an idea on the species’ dispersal abilities. The second type of hypothesis (hypothesis 2 and 3) was based on the species biology and the third type of hypothesis (4 and 5) was based on habitat connectivity in a large sense. In hipothesis 4 and 5 we considered that bocage landscape was difficult to cross because this element contained essentially agriculture zones with low vegetation in Ifen maps. In each one of these hypothesis the travel cost was fixed at 1 when we considered that the landscape features were easy to be crossed and it was fixed at 10000 for the features that we considered that they should be avoided by Pathmatrix algorithm because they were considered as too robust barriers to dispersion (Table 1).

**Table 1.** Traveling costs depending on landscape features

| Kind of habitat | Pixel traveling costs in each hypothesis |  |  |  |  |  |
| --- | --- | --- | --- | --- | --- | --- |
|  | IBD | Hypothesis 1 | Hypothesis 2 | Hypothesis 3 | Hypothesis 4 | Hypothesis 5 |
| Open space | 1 | 10000 | 10000 | 10000 | 10000 | 10000 |
| Coniferous forest | 1 | 10000 | 10000 | 1 | 1 | 1 |
| Bocage | 1 | 10000 | 1 | 1 | 10000 | 10000 |
| Deciduous forest | 1 | 1 | 1 | 1 | 10000 | 1 |

The specific hypothesis that were tested are presented here:

IBD (isolement by distance) : all the landscape features are considered as presenting the same travel cost. In this case the distances used are geografical (euclidian) distances.

Hypothesis 1: *Rhinolophus hipposideros* uses only decidious woods to move from one point to another because they are rich in insects and they allow him to hide from predators that are present in open spaces.

Hypothesis 2: *Rhinolophus hipposideros* uses preferentially the bocage and deciduous woods as dispersal corridors and he follows the same dispersal patterns that he displays when he is hunting at local scale avoiding open fields and coniferous woods poor in insects.

Hypothesis 3: *Rhinolophus hipposideros* uses his short and large wings in flight and has no difficulties to travel in open fields where travel distances and exposition time to predators are long.

Hypothesis 4: *Rhinolophus hipposideros* travels in coniferous forest that contribute to habitat connectivity.

Hypothesis5: *Rhinolophus hipposideros* travels in coniferous and deciduous forest that contribute to habitat connectivity.

We conducted Mantel tests in order to know if there was a relation between genetic and ecological distances using Fstat software (Goudet 1995). A Mantel test was also conducted to test the relation between genetic and Euclidian distances as it is usually applied to detect isolation by distance (Wright 1943). Finally, we conducted a last series of Mantel tests using the residuals of regression between ecological distances and genetic distances in order to eliminate the effects of euclidian distances.

### Analysis of the impact of landscape structure on genetic structure using an « a posteriori » method

« A posteriori » method consisted first in plotting genetic barriers with Monmonier algorithm implemented in Barriers 2.2 (Manni et al. 2004). Inference of genetic barriers is based on several steps. First of all we represent the colonies by georeferenced points in a map. We compute Delaunay triangulation in order to obtain a Vononoi tessellation. The Fst values corresponding to each pair of neighbor colonies are attributed to the corresponding sides of each triangle. The algorithm infers the location of the barriers following the decreasing values of Fst. As a consequence, we obtain barriers separing colonies that are genetically very different and regroup the colonies that are genetically close from each other. The robustness of the barriers was evaluated by bootstrap.

Finally, we look on the map if the locations of genetic barriers corresponded to the presence of particular landscape elements.

## Results

### Sampling

Applying logical CMR cumulating graphicals we obtained that the number of individuals by colony was comprised between 18 and 84. We present in **Table 2** the number of samplings and the number of individuals present.

**Table 2.**
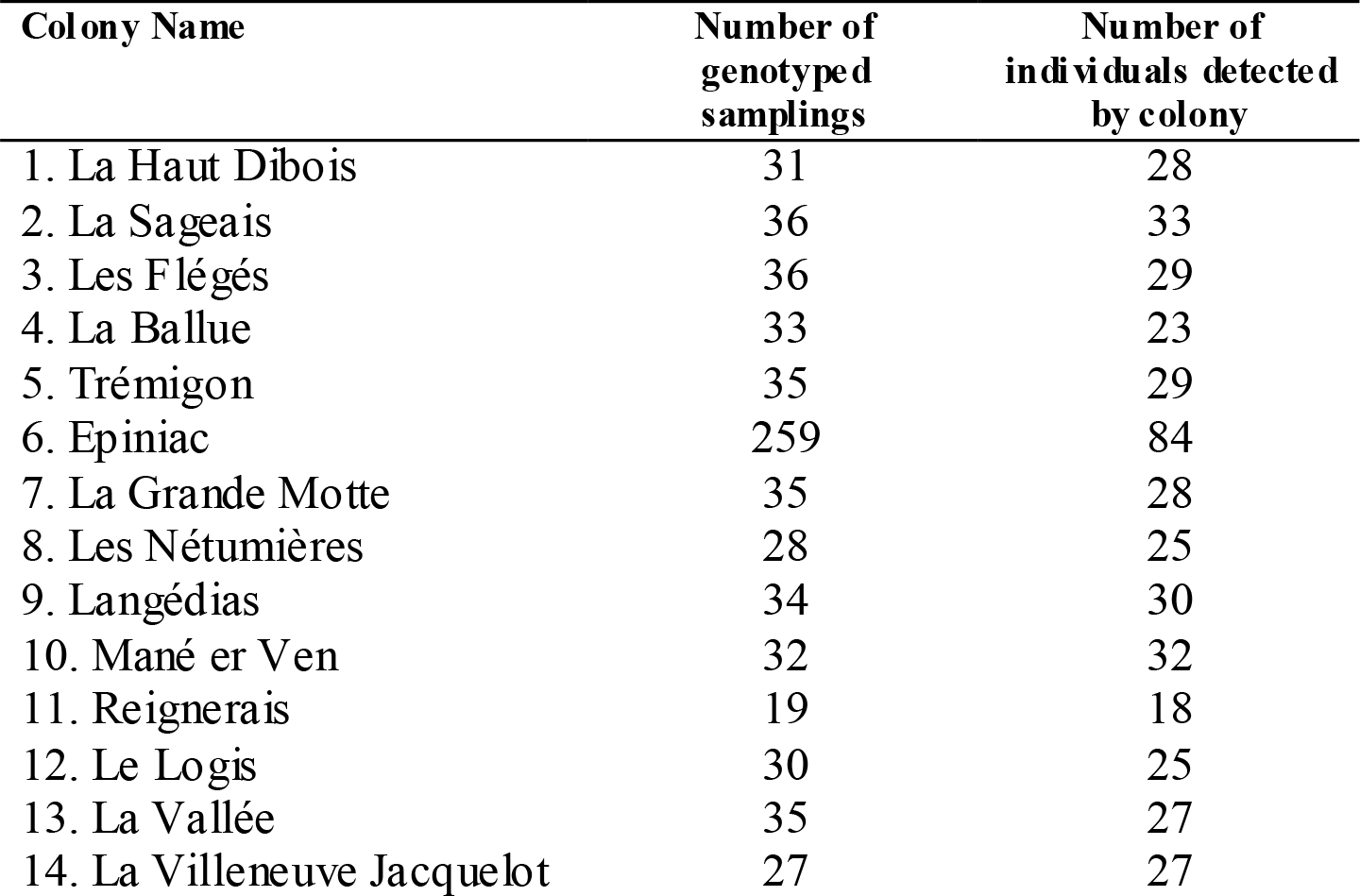
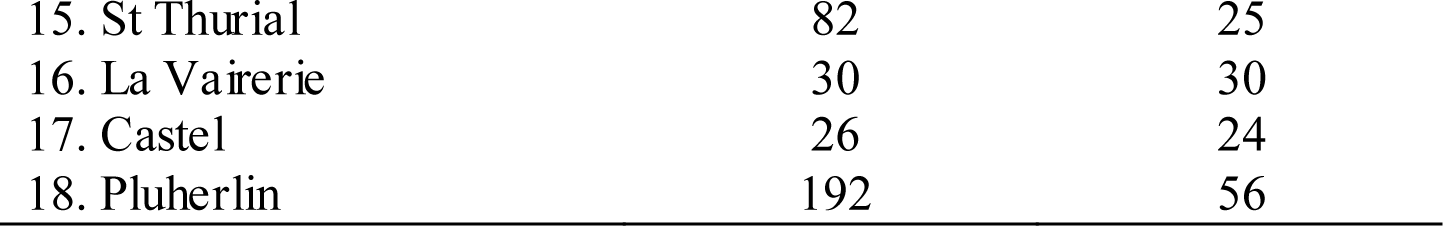
Total number of samplings studied by colony and number of detected individuals

### Genetic diversity and genetic distances

Preliminary tests conducted on the different colonies have shown that they were poorly differentiated but that this differentiation was significant (Fst = 0,057; 95% confidence interval: 0,050-0,065). Some colonies presented heterozygosity values inferior to the expected ones indicating that they are probably not at Hardy Weinberg equilibrium. When we look in detail what happened with each locus we detected that there was a genotyping problem with two loci: D2 and D119.

Moreover, a regression between Fst computed with and without taking into account the information brought by loci D2 et D119 has shown a deep modification of Fst values when we removed these two loci from the analysis (Fig. 1). By consequence, we decided to remove these two non reliable loci from all the analysis and keep only the six reliable loci.

**Figure 1.**
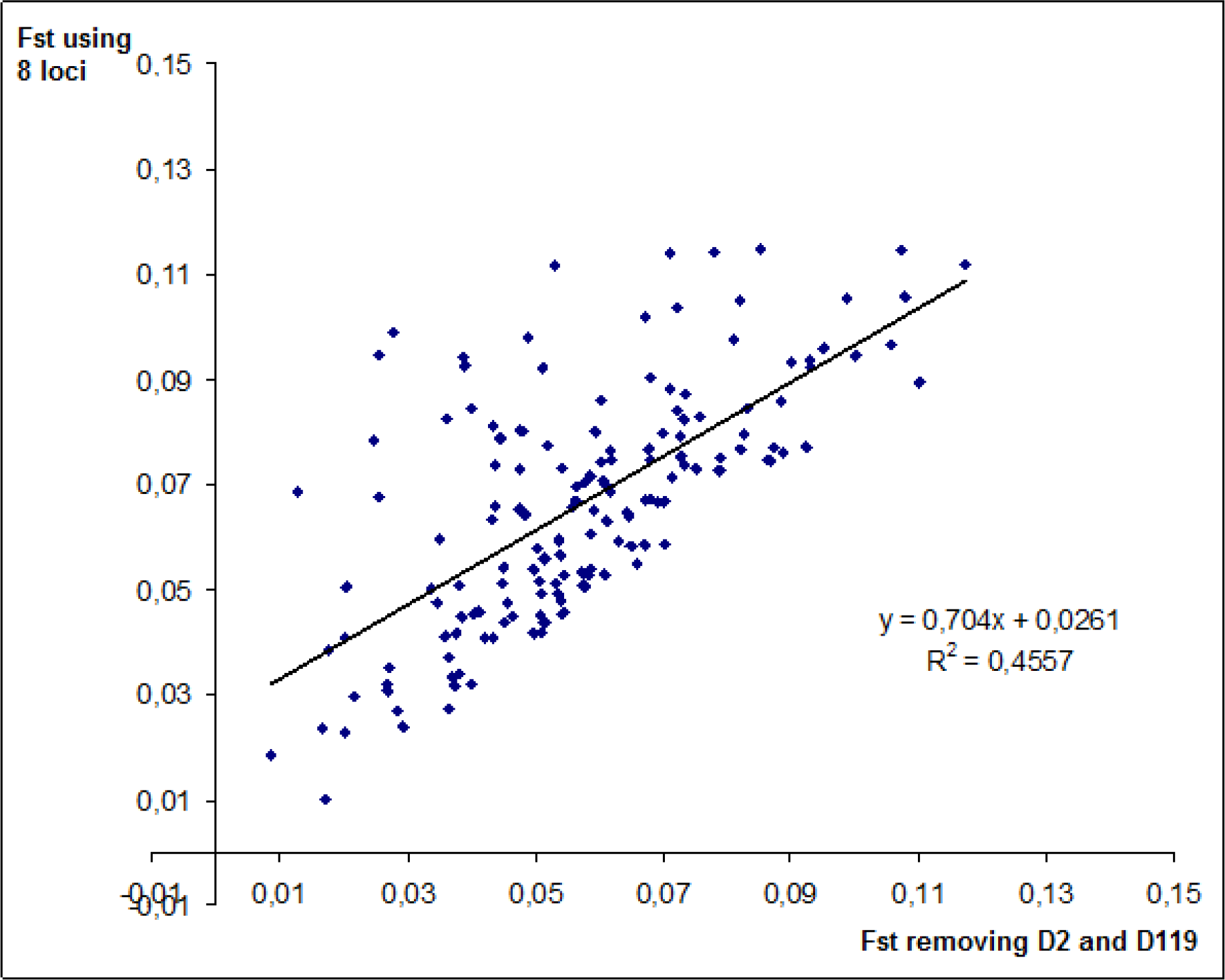
Linear regression of Fst colony pairs values using 8 and 6 markers

**Figure 2.**
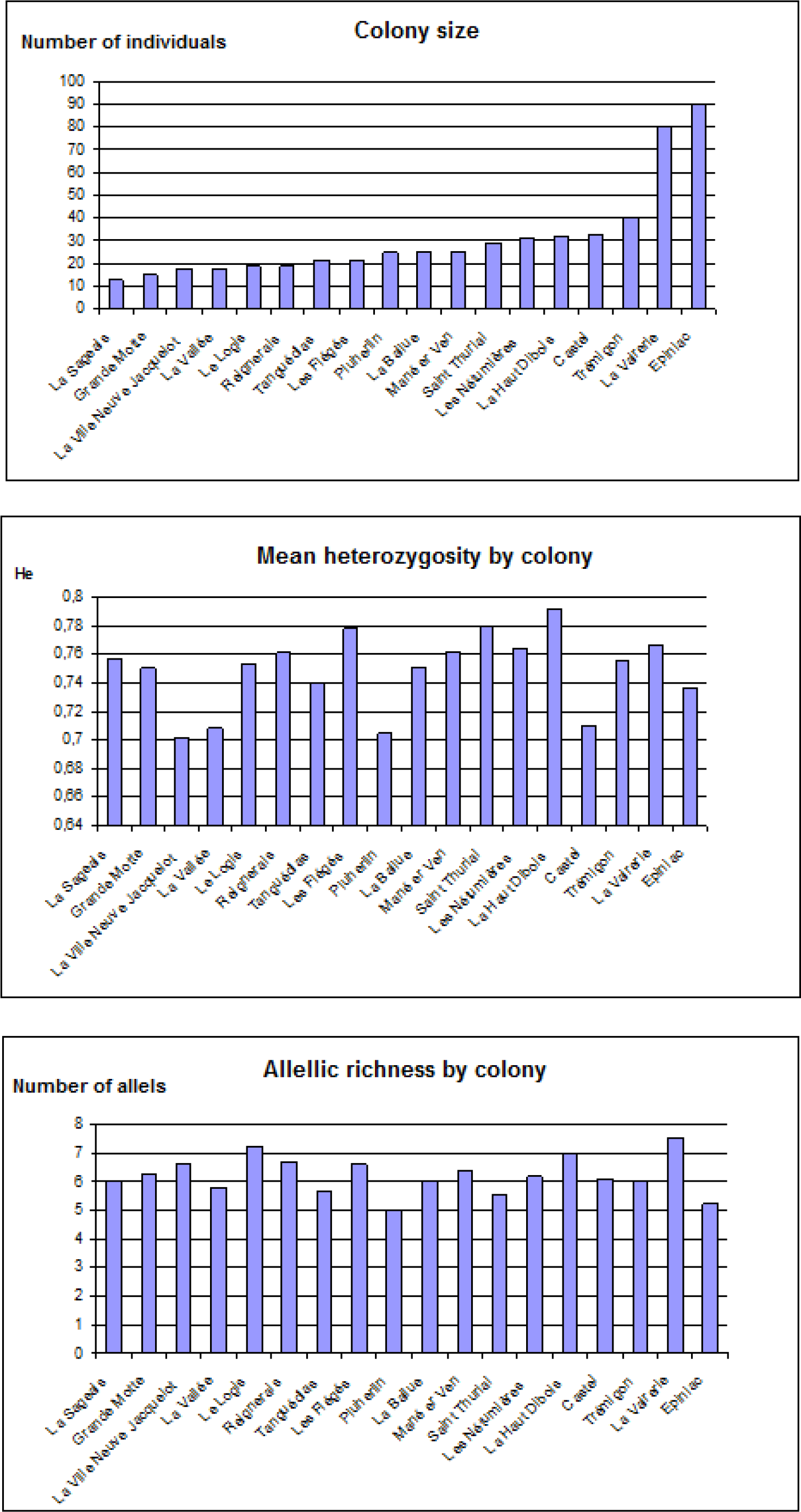
Size, Expected heterozygosity and allelic richness by colony

Differentiation tests between pairs of colonies have shown that they were all differentiated pair by pair (p<0,05) except Les Flégés and Nétumières colonies (Annex 1).

Concerning the 270 tests of disrupted link equilibrium conducted by pairs of loci, only 10 were significant after Bonferonni correction and they were distributed on different colonies and on different pairs of loci. By consequent we kept the whole information concerning the 18 colonies and the following loci: C108, D111, D9, D113, D102 et D103.

Allelic richness varied between 4,97 and 7,5 alleles per locus et per colony, and expected heteroygosity was comprised between entre 0,702 et 0,791. Colony size doesn’t seem to have a particular influence on these two parameters. Actually, we can observe small size colonies as Le Logis and Reignerais that present high values of heterozygosity and of allelic richness. On the contrary, some big colonies as Epiniac present low heterozygoties and allelic richness. Heterozygosity and allelic richness may be constrained by the degree of isolation of each colony. If we look the soil occupation map we can see that Pluherlin colonies, La Villeneuve Jacquelot and Castel are the most istant colonies from the group of colonies located in the northern east region. These three colonies present both low allelic richness and low values of heterozygosity.

This may be due to a low genetic flow and a low exchange of new alleles inducing isolation and a higher probability of consanguinity and low values of heterozygosity. On the map we can see that these colonies are surrounded by forests, the suitable conditions for colonies settlement are thus present but connectivity with other forest areas is probably not enough.

Colonies like Les Flégés, Le Logis, La Haut Dibois, Les Nétumières and Reignerais show the higher values of heterozygosity and allelic richness. They all belong to the areas were distances between colonies are weak. Genetic flow is probably facilitated by the proximity of colonies and by the presence of several forest areas observed on the map. Concerning this point it is important to underline that some individuals were detected by their droppings both in Les Flégés and La Ballue colonies, or in Les Flégés and Les Nétumières or in Reignerais and Languédias. The existence of these migrants brings information supporting the hypothesis that individual flow and genetic flow are probably more frequent in these areas. Languédias colony is interesting because it is less distant from the groups of the north and the east than the first studied colonies and it present intermediate allelic richness and heterozygosity values comprised between those of the northern-east and the southern-west. It is important to note that even if only 18 of the 50 Brittan colonies were sampled, the relative densities of colonies in the region were respected. Our remarks are thus in accordance with the field reality concerning the relative abundances of neighbors that each colony can have.

### Results of the « a priori » method

The « a priori » method allowed us to obtain different raster maps. We present two of them in the figure 3. The step maps used to compute them are presented in Annex 3.

**Figure 3:**
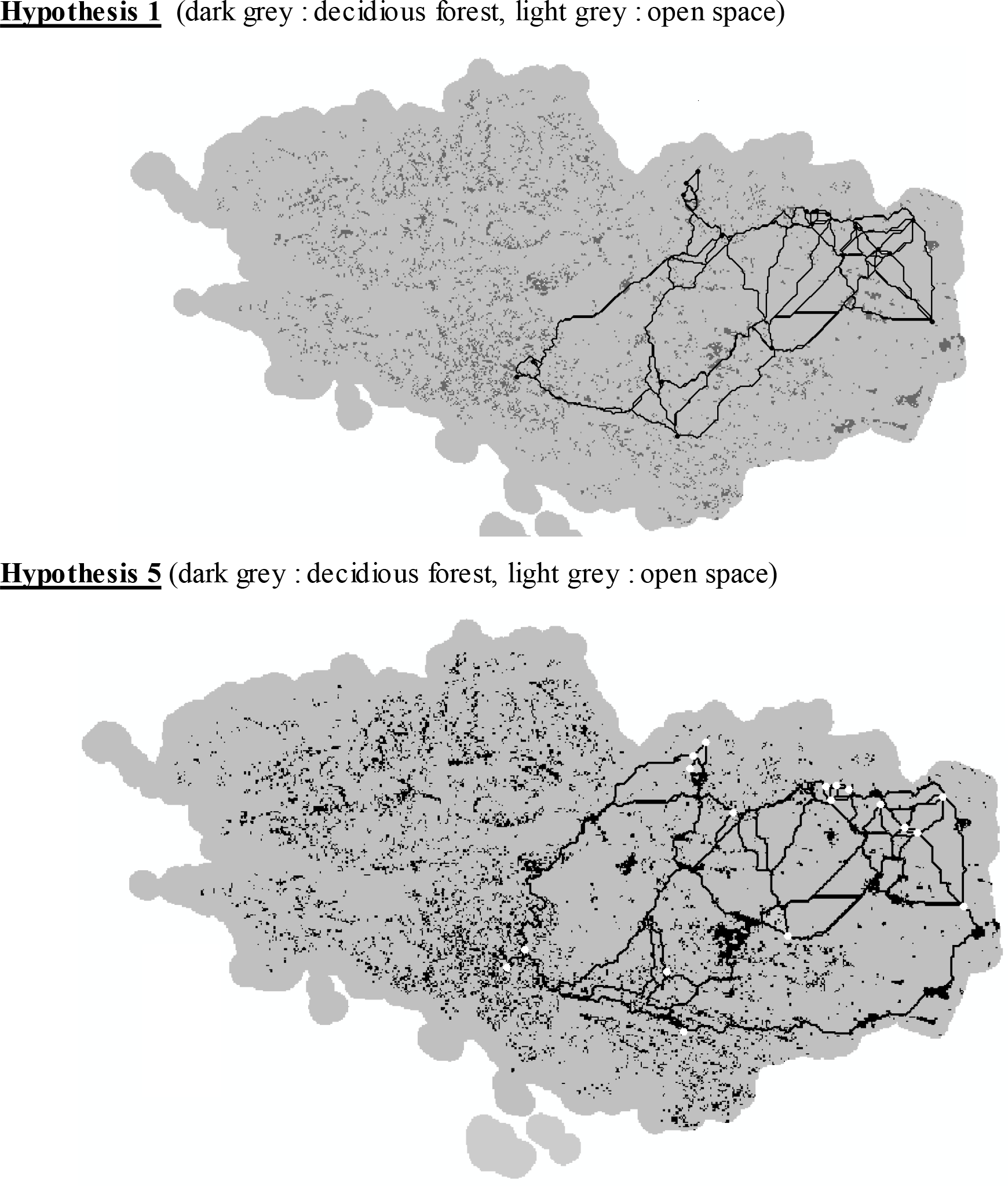
Paths obtained with Pathmatrix for hypothesis 1 and 5 (economic paths (1) in dark grey, costly pixels (10000) in light grey)

First of all, the results of Mantel tests revealed a significant correlation between the genetic distances and the Euclidian distances with a slope of 0,0072 which corresponds to a genetic isolation pattern that explains 6,07% of the genetic variance **Figure 4**. The first Mantel tests conducted to test the five hypotheses on habitat structure gave significant results too but they were very similar to those obtained when testing the isolation by distance hypothesis (Table 3). We thought that there was a strong correlation between ecological distances and euclidian distances so we decided to compute Mantel test between them. For the five hypotheses we obtained significant correlations and we saw that more than 98% of ecological distance value was explained by Euclidian distance.

**Figure 4.**
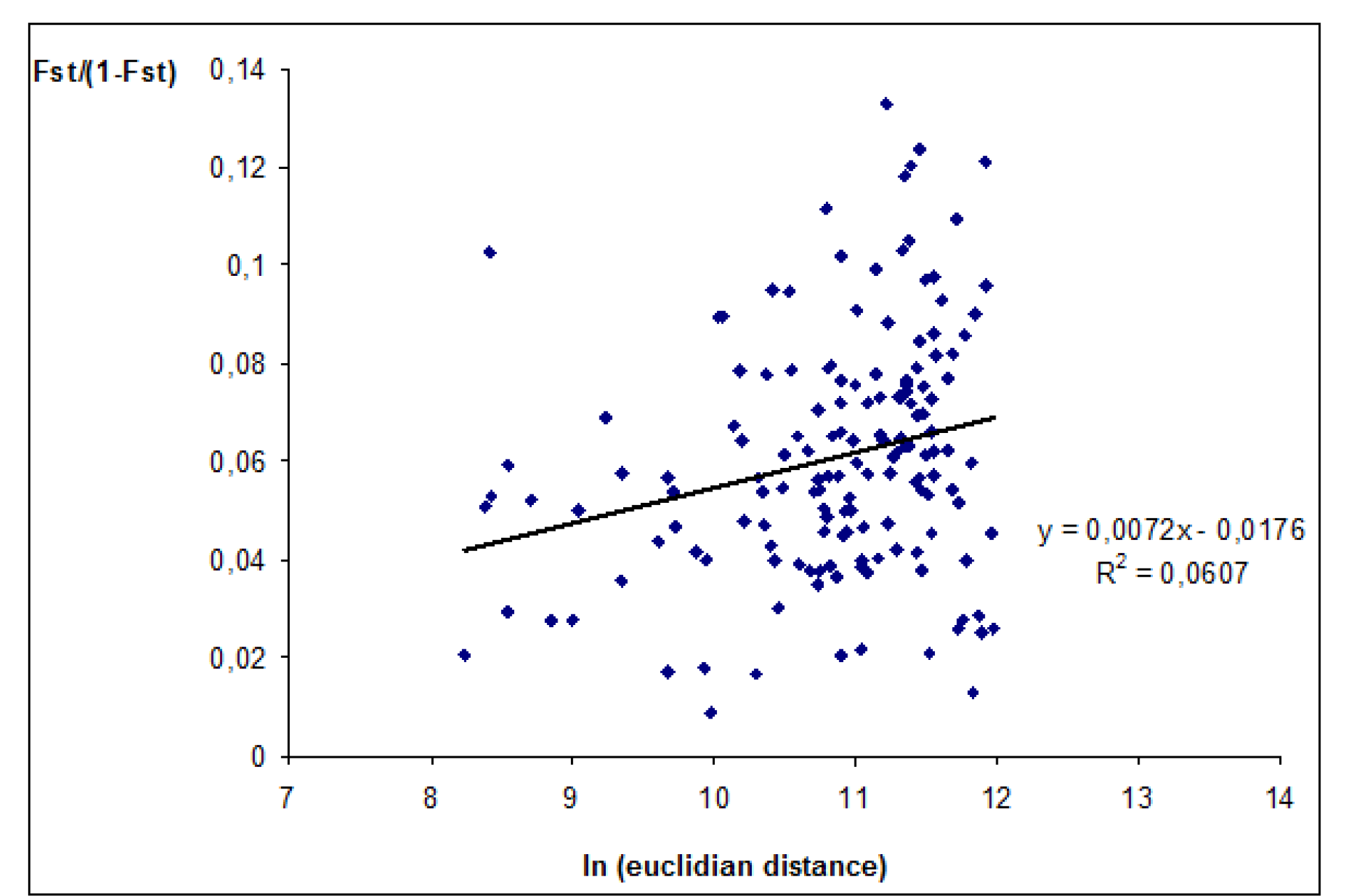
Isolement par la distance chez le Petit rhinolophe (*Rhinolophus hipposideros*)

**Table 3:** Percentage of genetic variance explained by each hipothesis and p values obtained in Mantel tests (Test 1: genetic distance vs ecological distance; Test 2: Residuals of regression between genetic distances and Euclidian distances vs residuals of regression between ecological distances and Euclidian distances

|  | % of variance explained by tests 1 | p values for tests 1 | % de variance explained by tests 2 | p values for tests 2 |
| --- | --- | --- | --- | --- |
| **IBD** | 6.07 | 0.001 | - | - |
| **Hypothesis 1** | 6.37 | 0.001 | 0.46 | 0.409 |
| **Hypothesis 2** | 6.43 | 0.005 | 0.60 | 0.360 |
| **Hypothesis 3** | 5.99 | 0.002 | 0 | 0.995 |
| **Hypothesis 4** | 6.17 | 0.001 | 0.11 | 0.689 |
| **Hypothesis 5** | 6.60 | 0.002 | 1.18 | 0.188 |

In order to avoid the redundant part of variance due to the euclidian distance, we conducted a second series of Mantel tests on the residuals of the latest regressions **Table 3**. The residuals of regression between genetic and euclidian distances were not significantly correlated with the residuals of the regression between ecological and euclidian distances p<0,05.

### Results of the « a posteriori » method

Thanks to Barriers 2.2 software (Manni et al. 2004) we could obtain the Delaunay triangulationbetween the 18 studied colonies. The inference of genetic barriers allowed us to distinguish the groups of colonies using Monmonier algorithm **Figure 5**.

**Figure 5.**
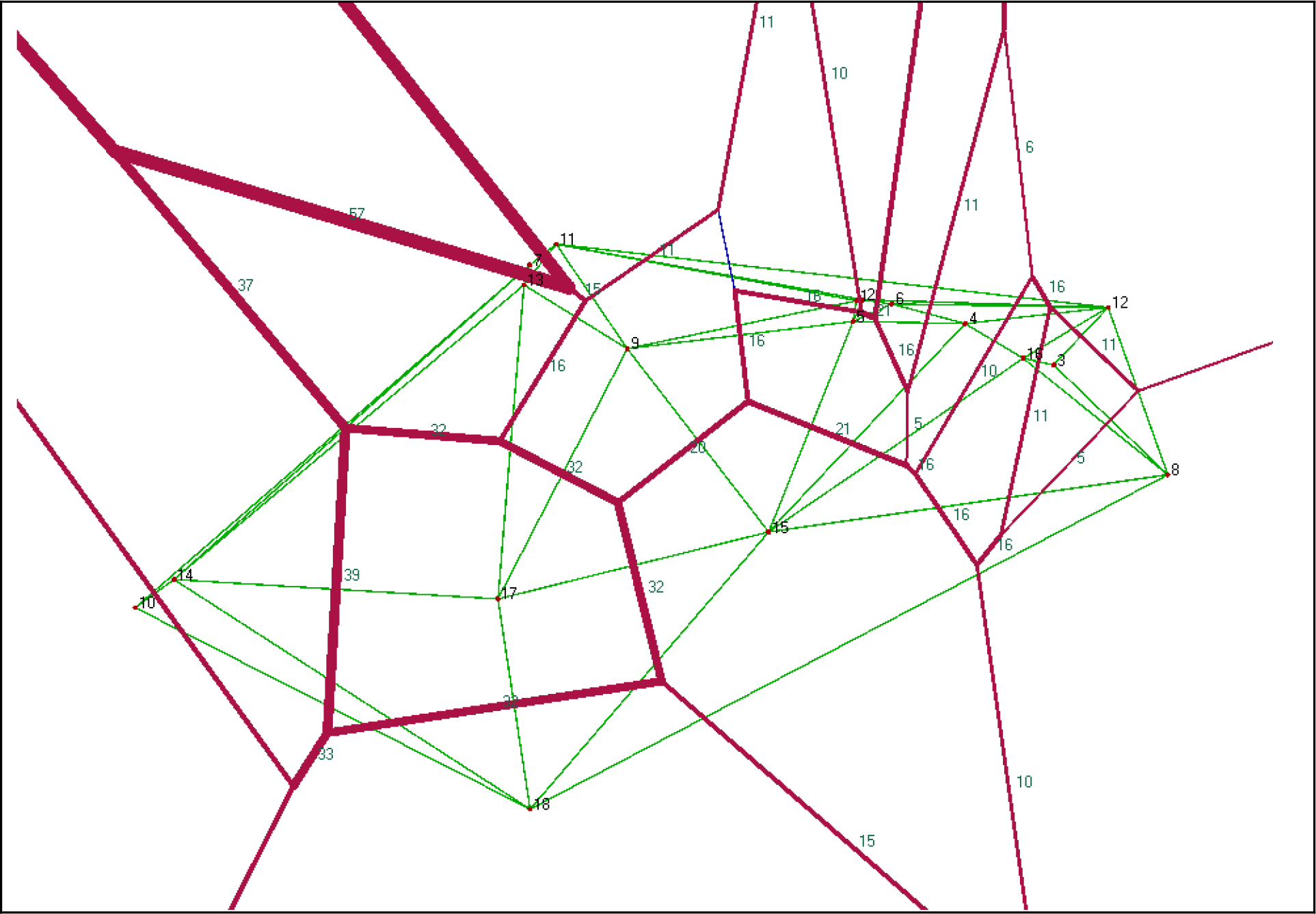
Genetic barriers obtained with Monmonier algorithm (colonies are represented by par the tops of triangles and their numbers correspond to those presented on Table 2). The numbers on the red barriers correspond to bootstrap values obtained for each one of them.

Most of the barriers are not very robust (bootstrap values < 58%). The more robust genetic barriers are located on the south-west side of the region. The importance of these barriers is due to isolation by distance which is stronger for the colonies of this area because they are more distant from the others. On the Corine Land Cover map from Ifen we notice that sometimes the barrier corresponds to the presence of open habitats, it is the case of barriers between the following colonies : La Villeneuve Jacquelot (14) and la Vallée (13), Castel (17) and Languédias (9), La Vallée (1) and La Grande Motte (7), La grande Motte (7) and Reignerais (11). However, the presence of barriers does not always correspond to the absence of plant covert. In the case where colonies are distant it is difficult to know the part corresponding to isolation by distance and the part corresponding to landscape.

## Discussion

### Importance of isolation by distance in Rhinolophus hipposideros

Our first conclusion from the study of the genetic structure of *Rhinolophus hipposideros* is that he presents isolation by distance in Brittany. This phenomenon explains 6,07% of the variation observed in the genetic data. This isolation by distance patterns seems important and is stronger than the one observed in other European chiropters as *Myotis nattereri* (Rivers et al. 2005) , Myotis bechteinii (Kerth et al. 2005) and *Nyctalus noctula* (Petit et Mayer 1999). Actually, the more the regression slope is big the more the isolation by distance is important and reflects low abilities of dispersion. Thus, concerning the three chiropters presented here, the one who presents the best abilities of dispersion is *Nyctalus noctula* with a null isolation by distance pattern **Table 4**.

**Table 4.** Importance of the isolation by distance in several species from different taxa. Here the regression slope is computed between the genetic distance and the logarithm of euclidian distances

| Species | Taxa | Regression slope ( genetic distance (Fst/1-Fst) / ln euclidian distance (m) |  |
| --- | --- | --- | --- |
| *Tetra tetrix* | Gallinacea | 0,0083 | (Caizergues et al. 2003) |
| *Helix aspersa* | Mollusque | 0,0237 | (Arnaud et al. 2003) |
| *Carabus solieri* | Coleoptera | 0,2857 | (Garnier et al. 2004) |
| *Arvicola terrestris* | Rodent | 0,00005 | (Berthier et al. 2005) |
| *Myotis bechsteinii* | Chiroptera | 0 en dessous de 100km<br>0,0125 au dessus de 100km | (Kerth et Petit 2005) |
| *Rhinolophus hipposideros* | Chiroptera | 0,0072 |  |
| *Nyctalus noctula* | Chiroptera | 0 | (Petit et Mayer 1999) |

In some species like Murin of Bechstein, *Myotis bechsteinii*, we can observe two different patterns of isolation by distance. Actually, isolation by distance is null between populations that are separated by less than 100km (Kerth and Petit 2005). For colonies separated by more than 100km the isolation by distance pattern is similar to the one observed in *Rhinolophus hipposideros*. We notice that *Rhinolopus hipposideros* present always an important isolation by distance even between populations separated by low distances. It is interesting to observe here that *Rhinolophus hipposideros* presents dispersal abilities comprised between a mollusk like *Helix aspersa* and those of a little rodent, *Arvicola terrestris*. The abilities of dispersion of the Lesser horseshoe bat are similar to the ones of a bird that flies very little, the Tétra lyre (*Tetrao tetrix)*. These aspects let us think that *Rhinolophus hipposideros* is limited in its movements (probably because of its little size) and by consequence the genetic flow would be more frequently between closed colonies. The species displays a dispersal pattern in Japanese steps as *Rhinolophus ferrumequinum* (a species phylogenetically closer to the *Rhinolophus hipposideros* (Kimura and Weiss 1964; Rossiteret al. 2000).

However; it is possible that isolation by distance is not simply the result of low abilities in dispersion but it can be due to sedentary patterns or to philopatric behaviors.

It could be interesting to study specific behaviors as reproductive ones that could explain our results. In several species of chiropters females are philopatric and present a close structure with few maternal lineages in the same group. This is the case of *Myotis bechteinii* (Kerth et al. 2000) and *Rhinolophus ferrumequinum* (Rossiter et al. 2005). Genetic flow depends on males dispersal abilities but also on females’ ones. Both sexes join each other in mating sites very far from the birth colonies. The travels conducted between birth colonies and mating sites explain the absence of isolation by distance in *Myotis bechteinii* in areas separated by more than 100km. It would be interesting to know if *Rhinolophus hipposideros* presents a similar social structure or not and if there are philopatric patterns that reinforces the isolation due to the low dispersal abilities.

### Landscape effect on Rhinolophus hipposideros dispersal

In this study we did not found evidence to asses that there is a relation between genetic structure of *Rhinolophus hipposideros* in Brittany and the big forests of the region. Actually, none of the hypothesis of the “a priori” method concerning landscape brought more information on genetic distance variance than isolation by distance. We did not observe clear differences in hypothesis including deciduous or coniferous forests. However, several field studies revealed that *Rhinolophus hipposideros* avoids coniferous woods and that they forage mostly in deciduous forest (Holzhaider et al. 2002). This behavior is related to cognitive abilities to distinguish the presence of conifers or deciduous trees by analyzing the different types of echoes reflected by these two very different types of trees (Fig. 6 ; Grunwald et al. 2004)

**Figure 6.**
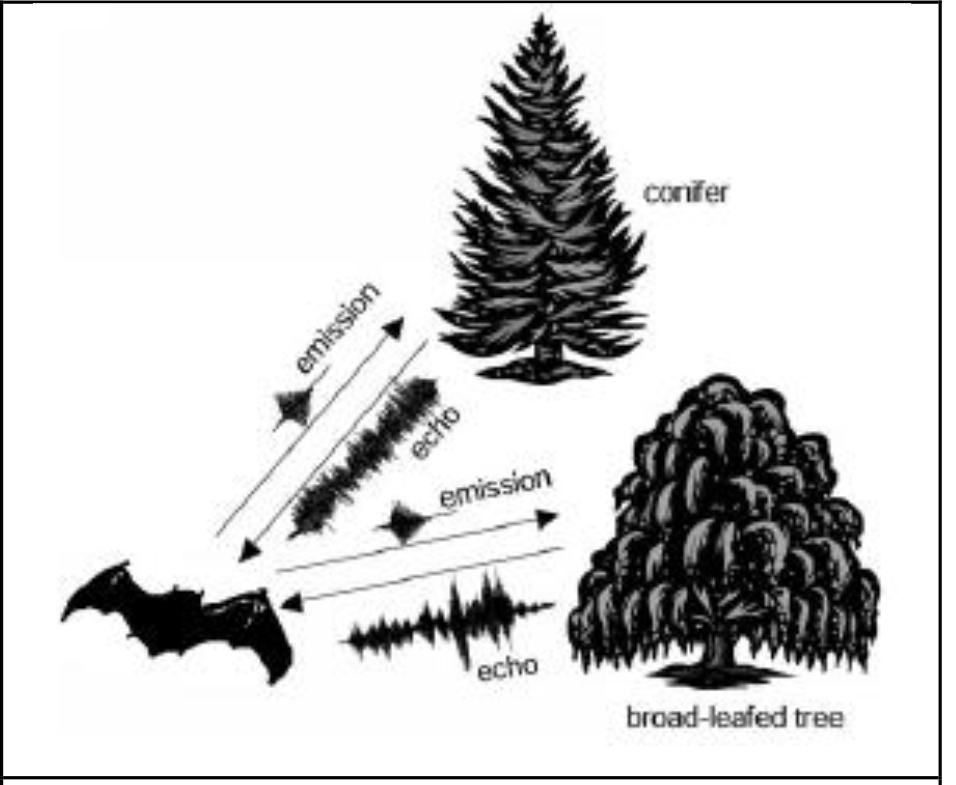
Different types of echoes reflected by different types of foliages. (Grunwald et al. 2004)

This aptitude to discriminate the two types of foliage is an advantage when taking decisions concerning the foraging sites as they have bigger probabilities to find insects when they hunt in deciduous woods (Tibbels and Kurta 2003). Tibbels and Kurta have demonstrated that deciduous wood areas which presented clearings and that were visited by chiropters *Myotis lucifugus*, *Myotis sodalist* and *Myotis septentrionalis* presented higher insects biomass than the other studied habitats. They found a high abundance of Dipters, Lepidopters and Trichopters (all these species that are included in *Rhinolophus hipposideros* diet).

The results we obtained with the « a priori » method seem to be contradictory with the ecological needs of the lesser horseshoe bat but it is not necessary the case. In fact, in this method we took into account the big forests and pixels used were of 25ha. Considering the behaviors that have been observed in other studies at other scales it is possible that the important landscape features in this bat dispersal are the hedges insuring connectivity and not the big isolated forests. In this order of ideas, if we combine the information brought by the « a posteriori » method and the information concerning hedges densities by Josso et al (1997) we found interesting clues for future studies on the impact of landscape on Rhinolophus hipposideros colonies. Actually, thanks to Monmonier algorithm we notice an ascendant gradient of robustness of barriers from the east to the west. In the south-west region we have a group of colonies separated by intermediate robust barriers while in the north-east region we have colonies that are separated by barriers that are twice weaker.

In their study « Les haies de Bretagne » Josso et al. (1997) represented the densities of different types of hedges on six maps (in m/ha). The six maps represented respectively: bocage embankments, low hedges, bushes, dominant bushes with big trees, big dominant trees with bushes, hedges composed only by big trees.

As Rhinolophus hipposideros flies preferentially between 2 and 5m over the ground, (McAney et Fairley 1988) we studied only the hedges comprising big trees, thus the three last maps. On these maps we notice that the north-east zone, previously mentioned is essentially covered by hedges which density is superior to 30m/ha and even superior to 65m/ha. En revanche, la zone située au sud ouest présente des densités de combinaisons futaies-taillis which densities are inferior to 10m/ha and very rarely some pixels in which the densities are superior to 30m/ha.

In the same way, the three colonies located at the northern top of the map, Reignerais (11), La Grande Motte (7) and La Vallée (13) are particularly interesting. These three colonies are very close from each other but they present very robust barriers genetic between. In this area, the hedge densities are inferior to 10m/ha. Or on constate que cette petite portion de la carte présente des densités de haies inférieures à 10m/ha. In conclusion, genetic barriers are more robust at the locations where hedges densities are lower.

It is important to remember that field studies conducted in the Finistère department have never found colonies in this area where hedges densities are very low. Josso and collaborators (1997) underline that the geographical distribution of colonies follows a clear gradient from the west to the east.

Bocage embankments at the north-west region are followed by dominant bushes in the central part of Finistère department. Then, big trees with bushes of the west central-region of Brittany and finally the big trees in the north region of Brittany. This opposition between bocage embankments at the west and big trees hedges at the east covers the distribution of decidious forests that are prevalent in the Finistère. The decidious big trees hedges are present mostly in Ille et Vilaine department.

### Perspectives

Ther are several pespectives for the study of the impact of habitat structure on populations’ genetic structure.

A more detailed analysis of the soil occupation maps and those of hedges densities as well as an analysis on the Pathmatrix way of computing distances could allow us to go further in the comprehension of eco-ethological processes that underline dispersal in Rhinolophus hipposideros.

Pathmatrix present the advantage of being able to compute the total cost of paths and to show which is the path globally less costly. However, even if this path is the less costly from one colony to another it doesn’t necessarily take into account the dispersal difficulties that un animal can be confronted with at a local scale. Fischer and Lindenmayer (2002), in their article on birds diversity talk about the SLOSS debate in which they ask the question to know what is more interesting in conservation biology between a big patch or several little patches. Here, we could think about this same question applying it to our particular context. It is possible that a big forest facilitates the dispersion of Rhinolophus hipposideros and that paths that cross them present lower total costs. However, if open spaces are also big, it would probably be more interesting to follow a series of patches close from each other and less costly at local scale even if at big scale it is more costly (fig. 7). At a local scale, it seems to be more important that the distance of an open space should not be too high.

**Figure 7.**
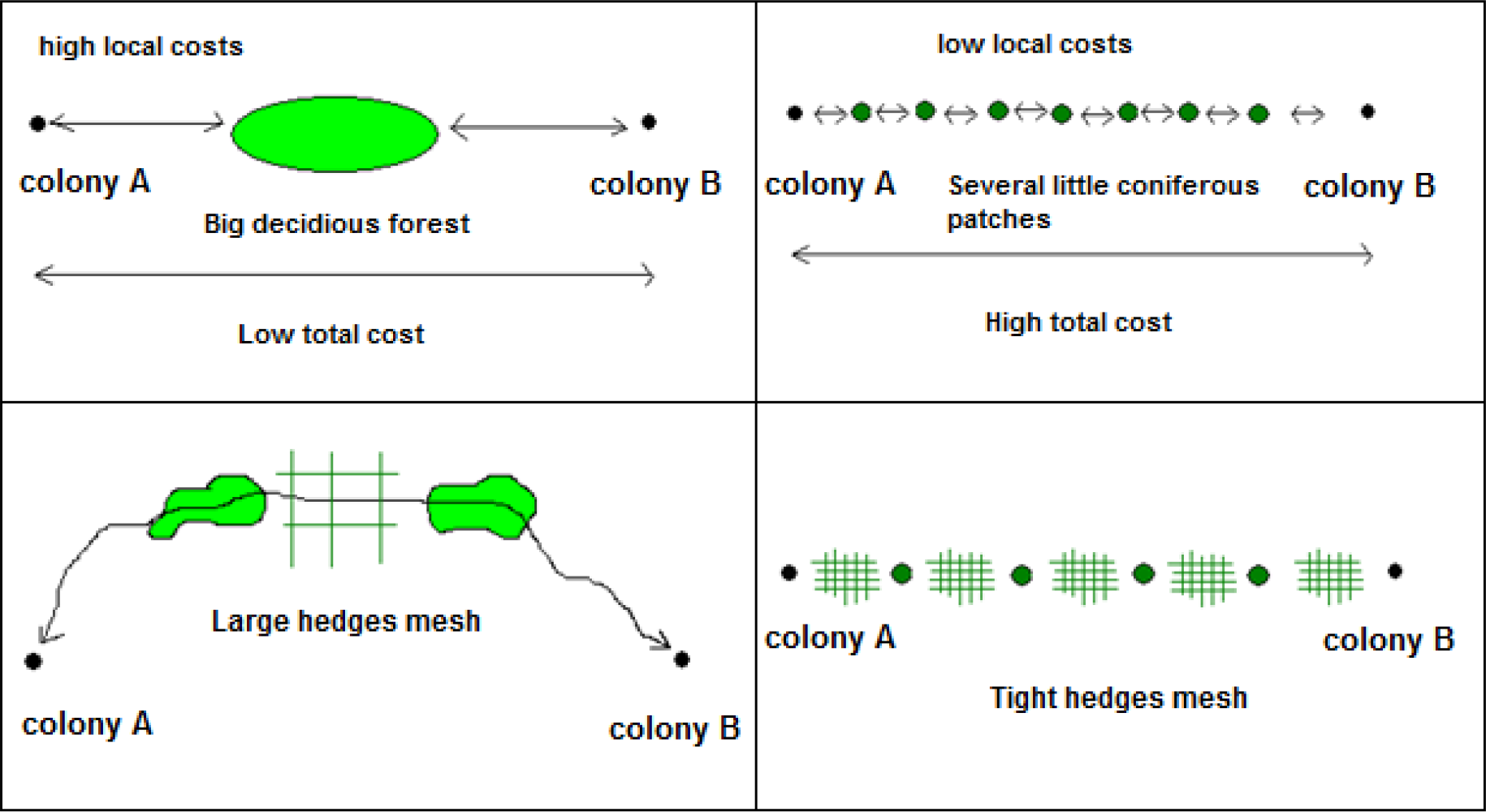
What distances are accessible and manageable for the bats ?

It would be very informative to analyze in a more precisely approach the distribution of the different patches and the hedges densities that exist between them (fig.8). In order to go further than we went in this study it would be interesting to conduct an “a priori” method using pixels reflecting hedges densities. It was not possible for us to compute this kind of mehod because we did not have such precise information. More studies and maps on soil use need to be done by the authorities that have the access to this kind of information.

In conclusion, when landscape preferences would be defined it could be also interesting if there are several species competing for the use of these preferendums and to know if there are or if there are not predators associated to these places. Arlettaz et al. (1999) noticed that the decreasing evolution of Rhinolophus hipposideros was coupled with an increase of *Pipistrellus pipistrellus* populations. They observed that the diets of these two species were very similar and that Pipistrellus pipistrellus catches more and bigger preys than Rhinolophus hipposideros.

It is necessary to study deeper the biology of the species and their interactions with their surrounding habitat in order to understand better their evolution and to be able to condut pertinent conservation projects.

---

**Annex 1.**
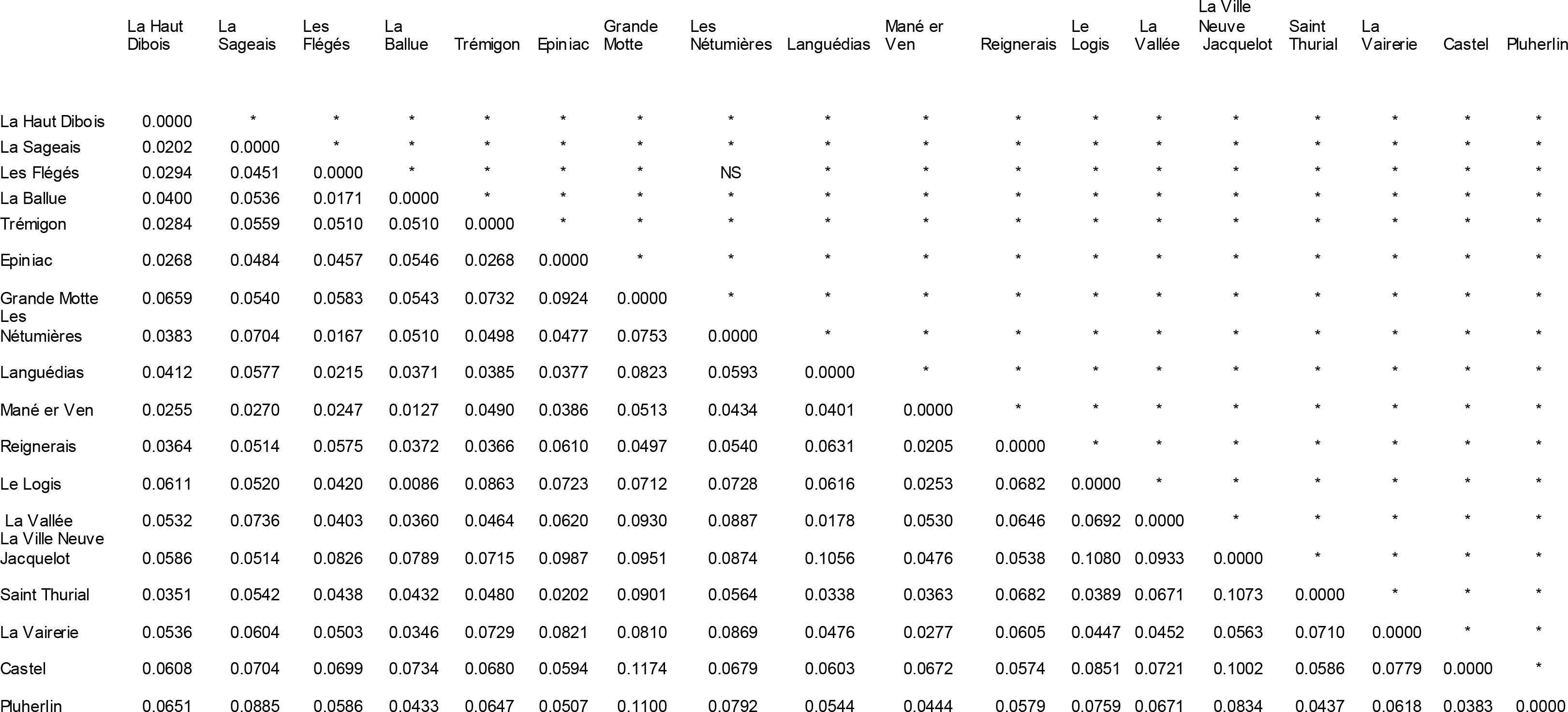
Fst significativity by pairs of colonies

**Annex 2.**
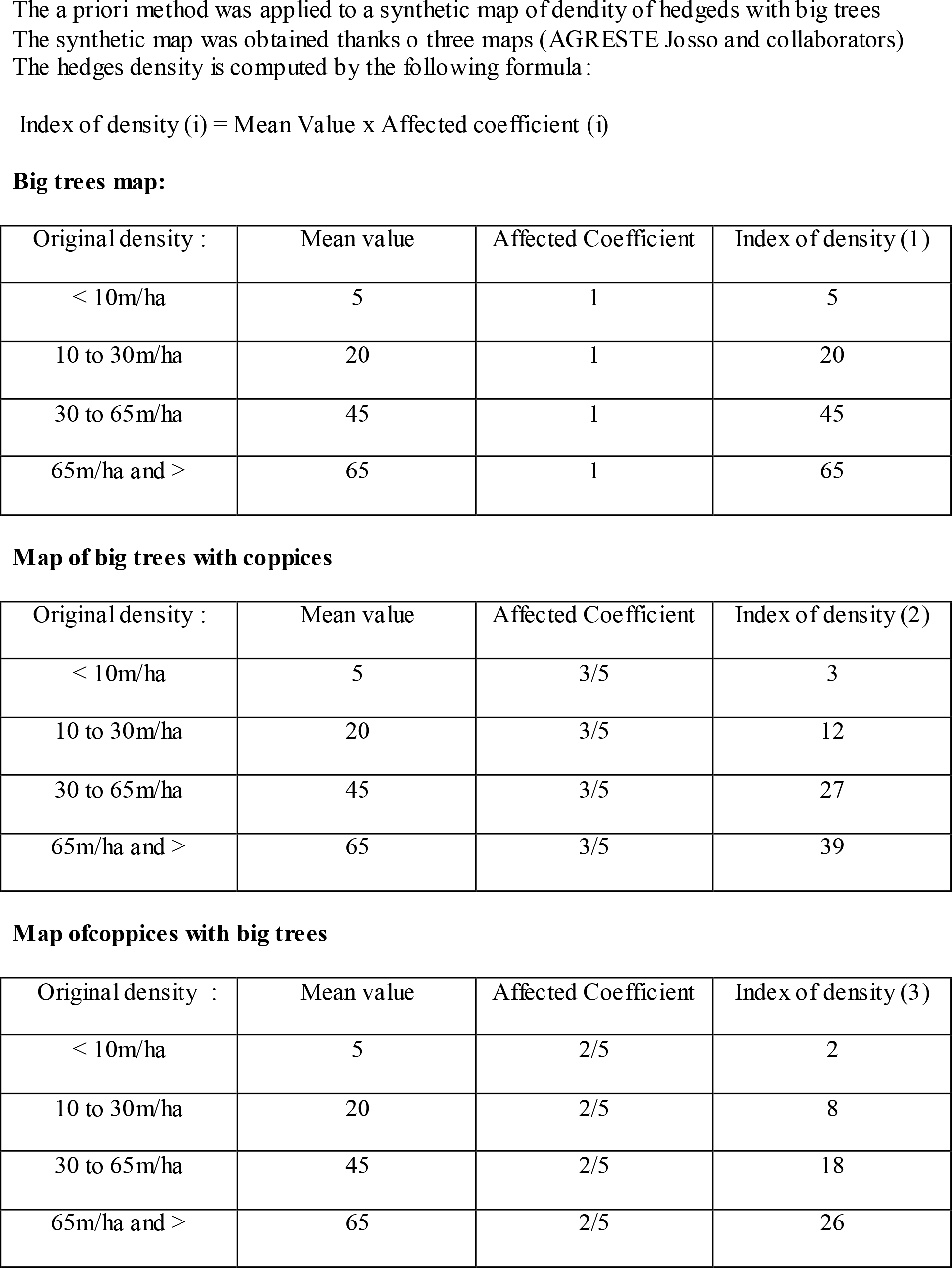
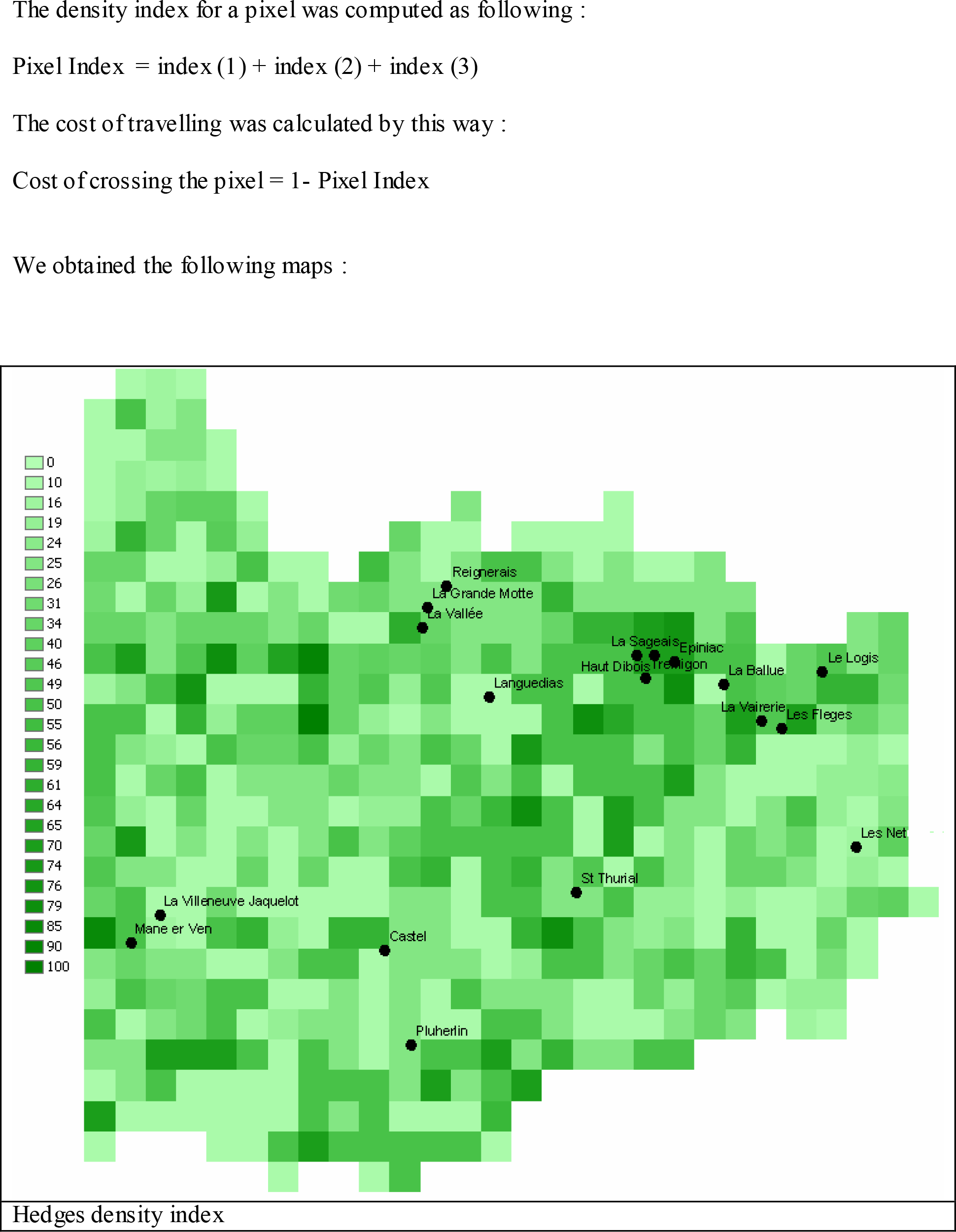
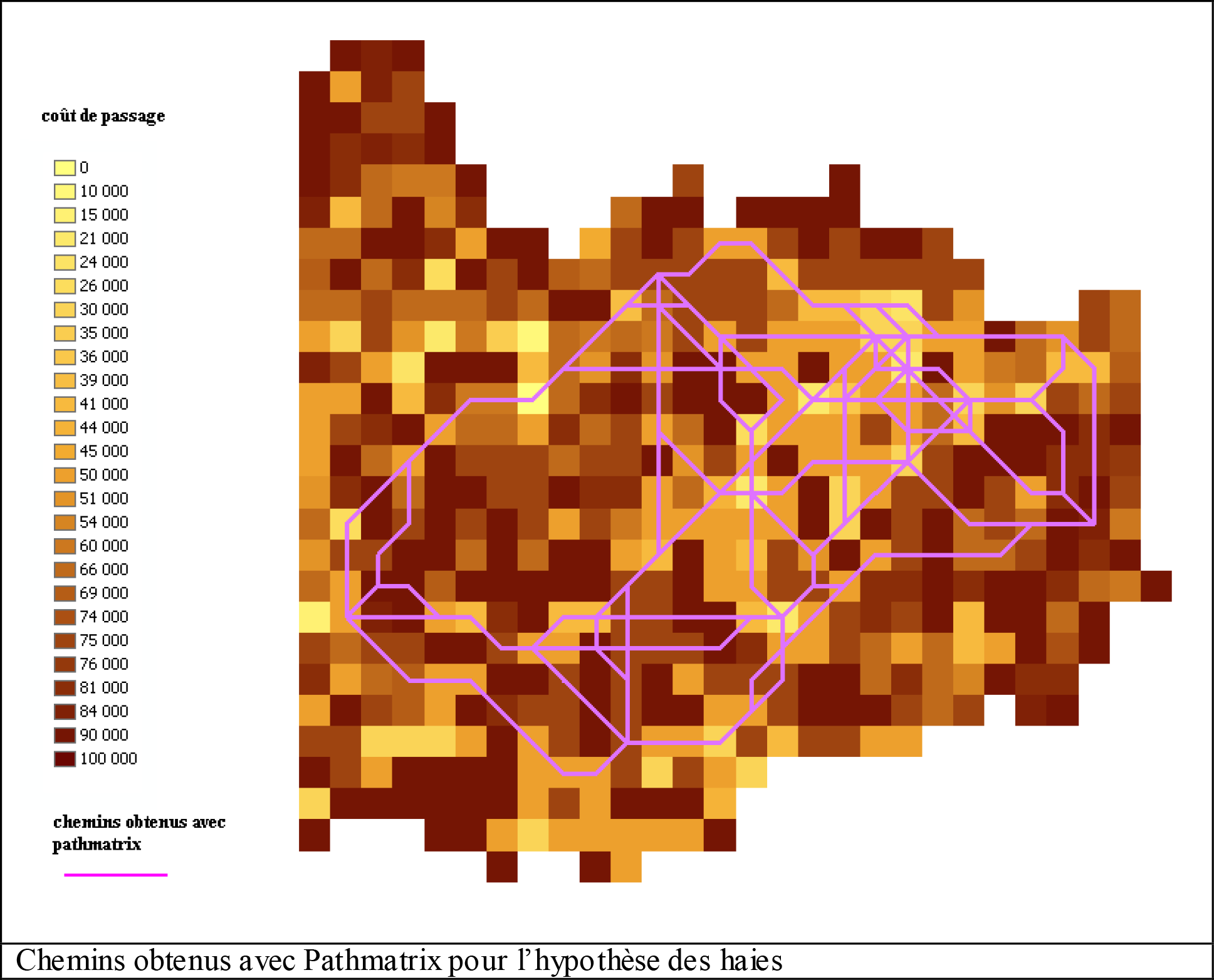
A priori method applied to hedges

**Annex 3.**
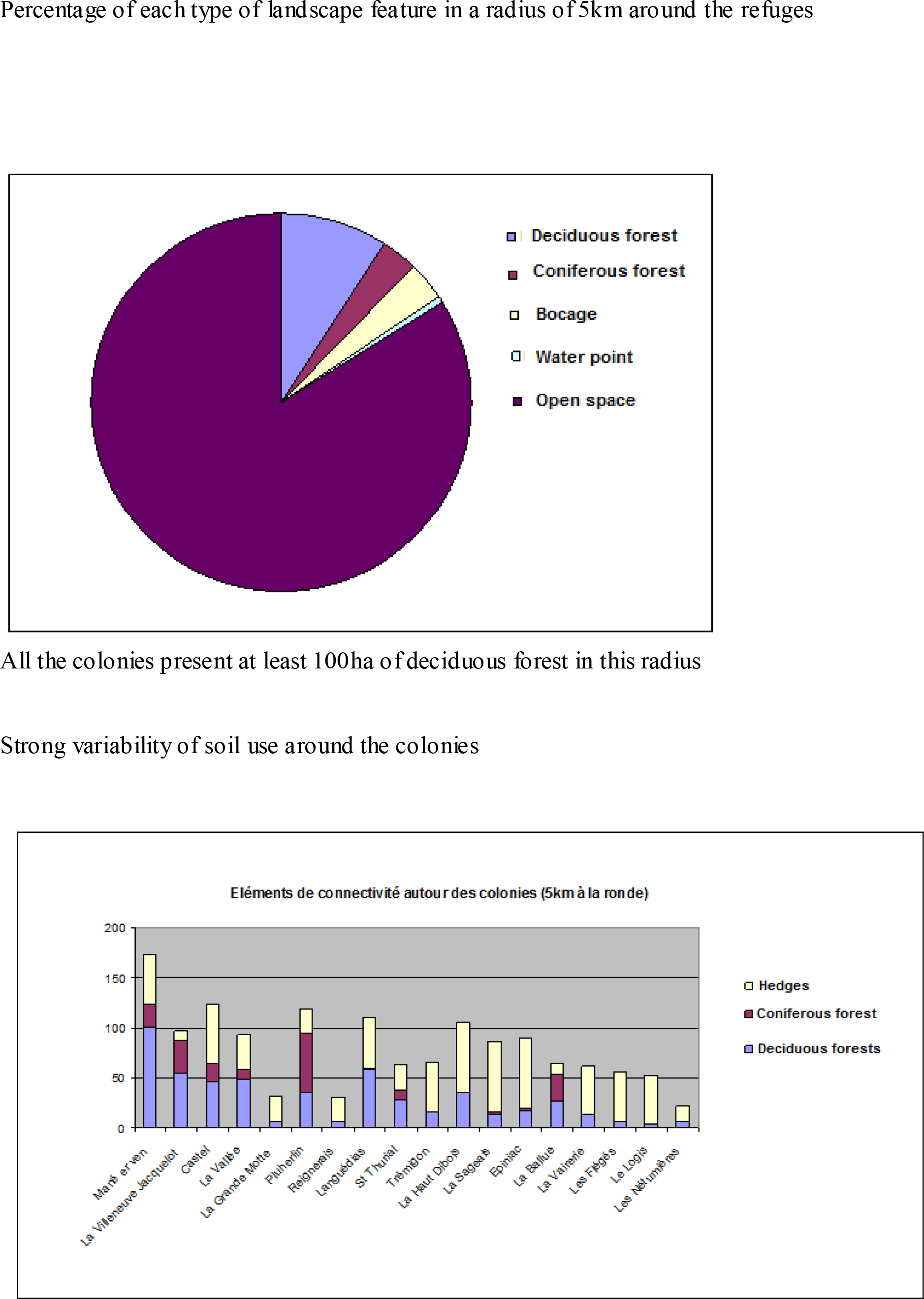
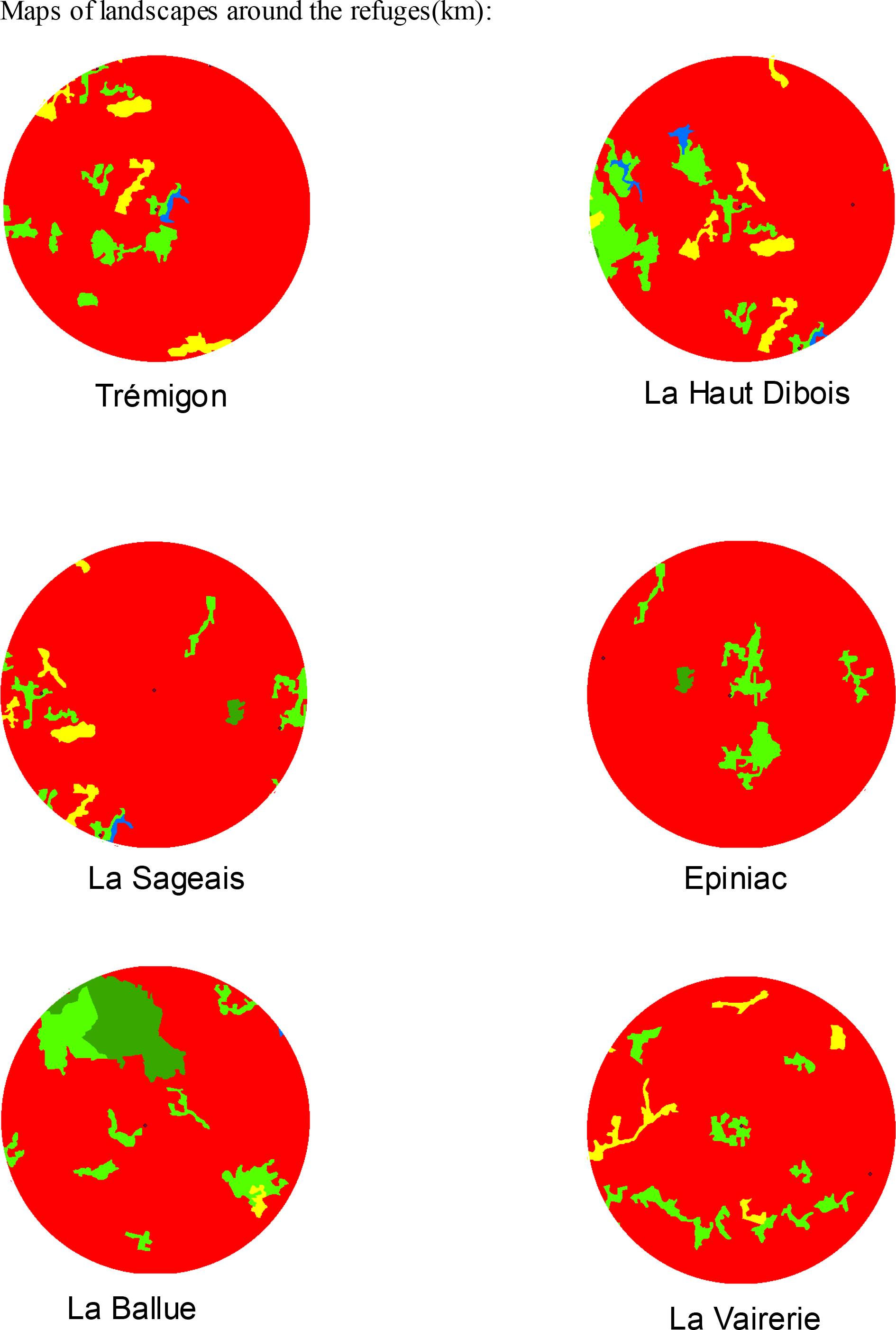
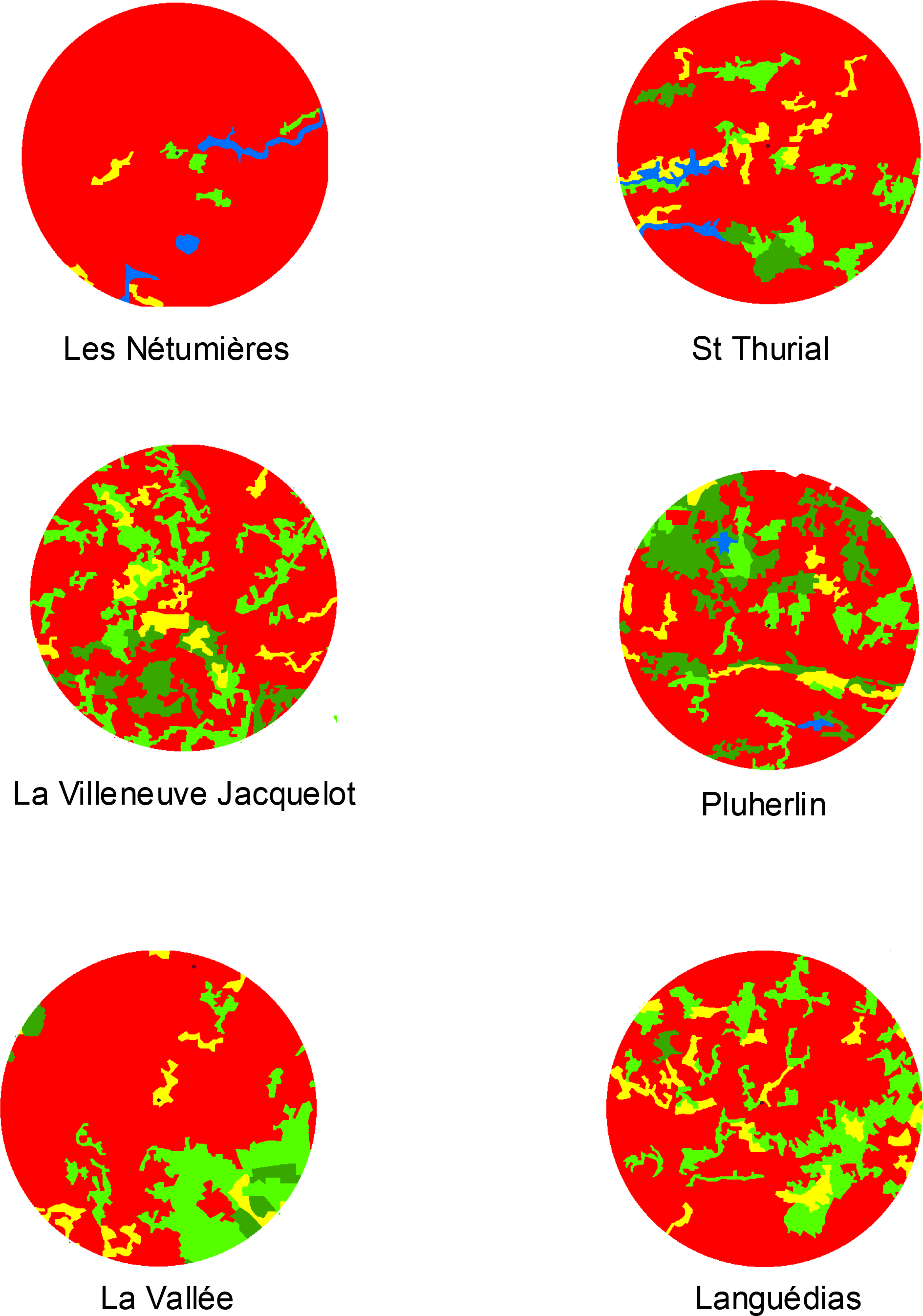
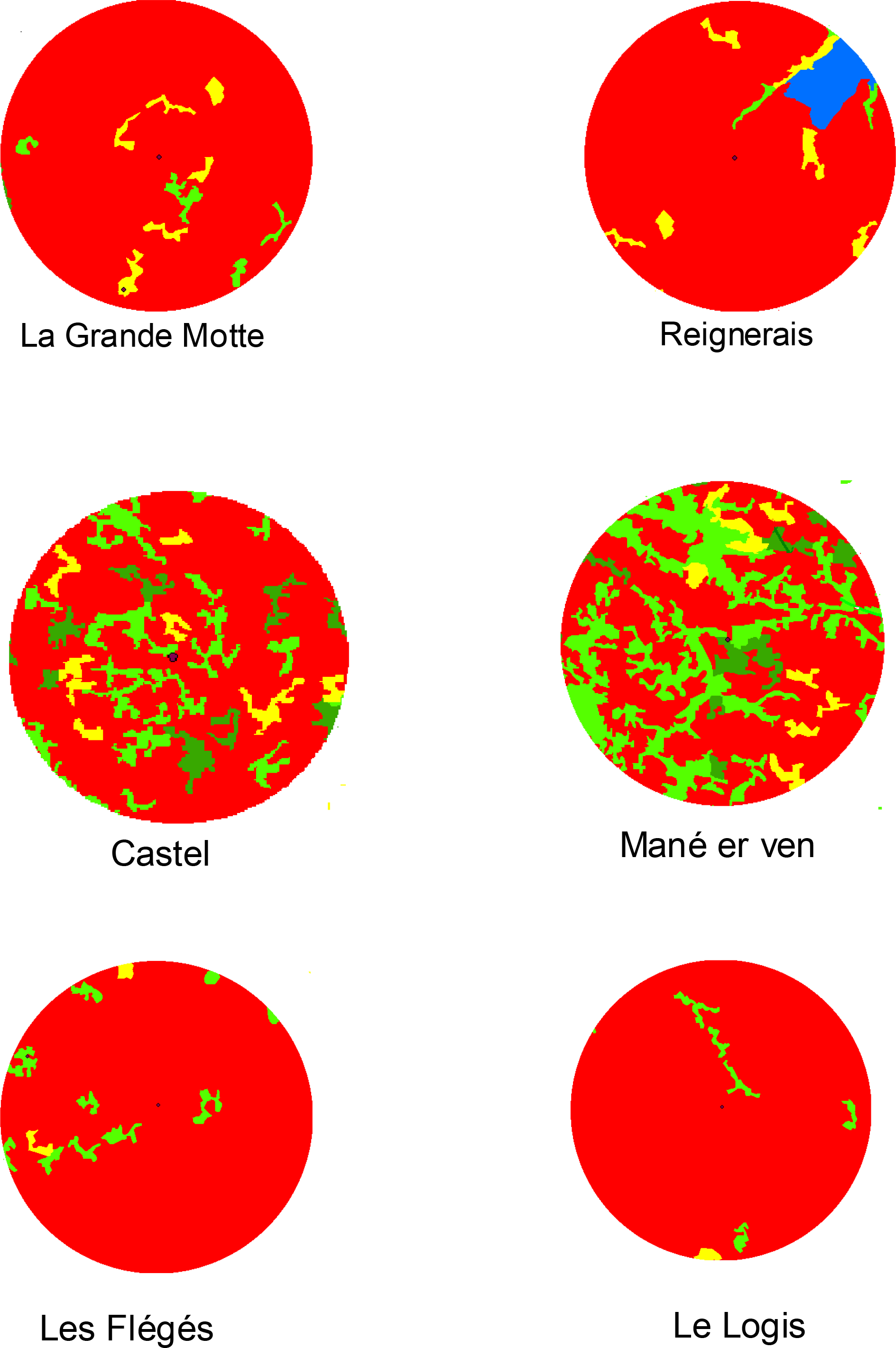
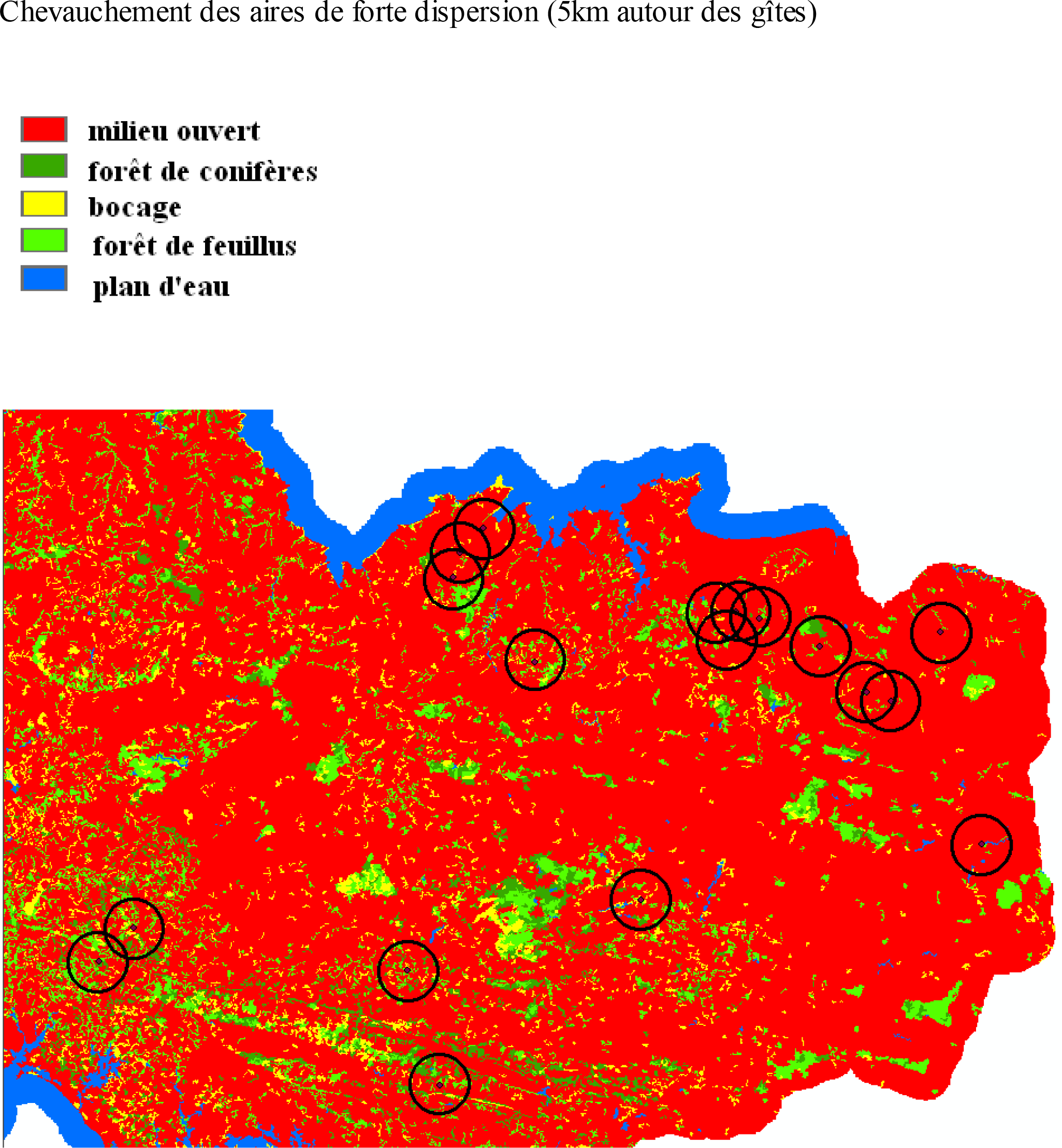
Landscape analysis around the refuges

